# Benchmark of Wide Range of Pairwise Distance Metrics for Automated Classification of Mouse Mutant Phenotypes from Flow Cytometry Data

**DOI:** 10.1101/2025.01.06.631468

**Authors:** M. May, T. Hewitt, B. Mashford, D. Hammill, A. Davies, T. D. Andrews

## Abstract

Precision medicine requires a comprehensive mapping of genotype to phenotype to provide patients with individually tailored treatment. However, when using flow cytometry to identify phenotypes, such as the quantity of various immune cell populations in tissue and blood used to identify autoimmune disorders, it is often unclear which cellular phenotypes are from healthy and disease individuals, especially when including the effects of population diversity, due to the high-dimensional nature of the data. To identify and segregate healthy phenotype from various disease phenotypes, we use pairwise distance metrics between each sample’s cell populations. By comparing distance metrics between C57BL/6 clone mice with mutations of known phenotype, we find that cosine similarity is best suited for segregating wildtype from mutant samples while respecting minute differences in already small cell populations, and that standardised Euclidean distance is best suited for machine-learning input due to its sensitivity. Both metrics outperform other tested metrics (including Aitchison, Euclidean, Manhattan, Earth-Movers Distance, and squared Euclidean). We demonstrate the utility of these different pairwise metrics through their application to a classification task of known mutant phenotypes: using an existing FACS phenotype dataset derived from X000 inbred C57BL/6 mice that harbour potentially phenotypic genetic variation introduced through ENU mutagenesis of individual pedigree-founding G0 male mice.

## 1 Introduction

Understanding the function of genes, genetic variation, and their phenotypic effect in individual people is an integral part of precision medicine (1). Precision medicine utilises knowledge of genomic variation to deliver treatments that are tailored to the individual, a method with the potential to greatly improve treatment outcomes (2). However, only genotypes with known phenotypes can be used in therapeutic targeting. Measuring the functional effect of mutation is therefore important to precision medicine, and careful attention must be paid to how functional effect is measured in order to avoid false conclusions (3, 4). Not only is a sensitive phenotyping method required, such as flow cytometry, but the ability to differentiate between biologically distinct populations, such as wildtype and mutant, is essential (3, 5).

Flow cytometry is a technology that, despite being over a century old, remains highly relevant and useful for the phenotyping task of quantifying each individual cell’s surface markers (6, 7). Fluorescent antibodies attach to cellular markers, allowing for up to thirty parameters (markers) in millions of cells in a sample at a time (8). In analytical flow cytometry, the parameters are then used to manually subdivide samples into cell populations (9), describing the quantity of varying cell populations in each sample.

Distinguishing between samples requires a measure that can cluster sample phenotypes based on a distance that represents biological reality. One such measure is a pairwise distance metric. Many distance metrics exist, each providing a different way to measure the difference between samples, and each having its own application it is suited towards (10, 11). Flow cytometry phenotypes are high-dimensional, which is a challenge due to the curse of dimensionality (12). Therefore, a systematic comparison of distance metrics is necessary to ensure biological validity is being captured instead of spurious correlations.

This paper identifies which distance metric best captures differences in cell populations between mice with wildtype and mutant phenotypes, then applies that information towards classifying immunophenotypes to group mice with others with a similar phenotype. A set of distance metrics are appraised based on their ability to distinguish between wildtype and mutant mouse immunophenotypes, and the magnitude of that distance. We identify distance metrics that are best suited to this task and make recommendations for future research on which metrics are biologically relevant.

## 2 Results

### Input Datasets

Two datasets of mice were used: the first dataset (dataset A) was designed to identify the distance metric most reflective of biological reality by using CRISPR to induce specific known mutations in C57BL/6 mice. It consisted of spleen samples collected from five C57BL/6 mice: three wildtype clones and two mutants, Kenobi (C57BL/6JSfdAnu) and Rag KO (Figure 1A). The data has been compensated, cleaned of cellular debris and singletons, and cells were manually segregated into cell populations (gates) through a series of scatterplots comparing two markers. These five mice were spread over three plates (each plate with a different quantity of antibodies) and two technical repeats. The plates share a common set of 12 markers (B220, CD3, CD8, CD19, CD25, CD44, CD62L, IgD, IgM, Ly6C, NK1, and LD).

The second dataset (dataset B) was the product of chemical mutagenesis experiments utilising *N*-ethyl-*N*-nitrosourea (ENU). It is noisier and was used to test the reliability of the distance metrics under challenging conditions. It consisted of sixty G3 mice that were part of an ENU mutagenesis experiment. As above, the data has been compensated, cleaned of cellular debris and singletons, and gated. These mice were run on the same plate and share a common set of eight markers (B220, CD3, CD4, CD44, IgD, IgM, KLRG1, and NK1).

### Distance Metrics

Of distance metrics used to compare immunophenotypes, cosine similarity and standardised Euclidean distance were the most accurate in differentiating wildtype to mutant immunophenotype. As dataset A consisted of known disease phenotypes caused by known mutations, it was used as a baseline to determine which distance metrics reflected biological validity. We evaluate distance metrics by observing how each measures the distance between groups of mice with differing immunophenotypes.

A biologically meaningful distance metric should show progressively larger differences between mouse groups that are increasingly distinct, rather than simply reflecting technical variability or background noise. First, we should expect the lowest distance should be between technical replicates. Second, a low distance between the three wildtype mice, as we expect mostly complete uniformity amongst them because they are clones. Third, an intermediate distance between the two kinds of mutant mice. Fourth, the highest distance should be between wildtype and both kinds of mutant mice.

### Choice of Distance Metric Determines Segregation of Immunophenotype that is either Resistant to Noise (Cosine) or Sensitive to Individuals (standardised Euclidean)

When observing the difference in absolute cell count in gates between samples in dataset A (Figure1B), wildtype samples are all similar to each other while Kenobi and Rag are both very different to wildtype and to each other.

As all distance metrics showed progressively larger differences between increasingly distinct mouse groups, all distance metrics tested were showed some level of reflecting biological reality and being robust to instrumental noise.

Distance metrics were either robust to differences between individuals and technical replicates or sensitive to them. Figure 2A observes that cosine similarity, correlation, and squared Euclidean finds a small difference between wildtype samples, and a relatively very high difference between mutant samples and wildtype-to-mutant samples.

**Figure 1.**
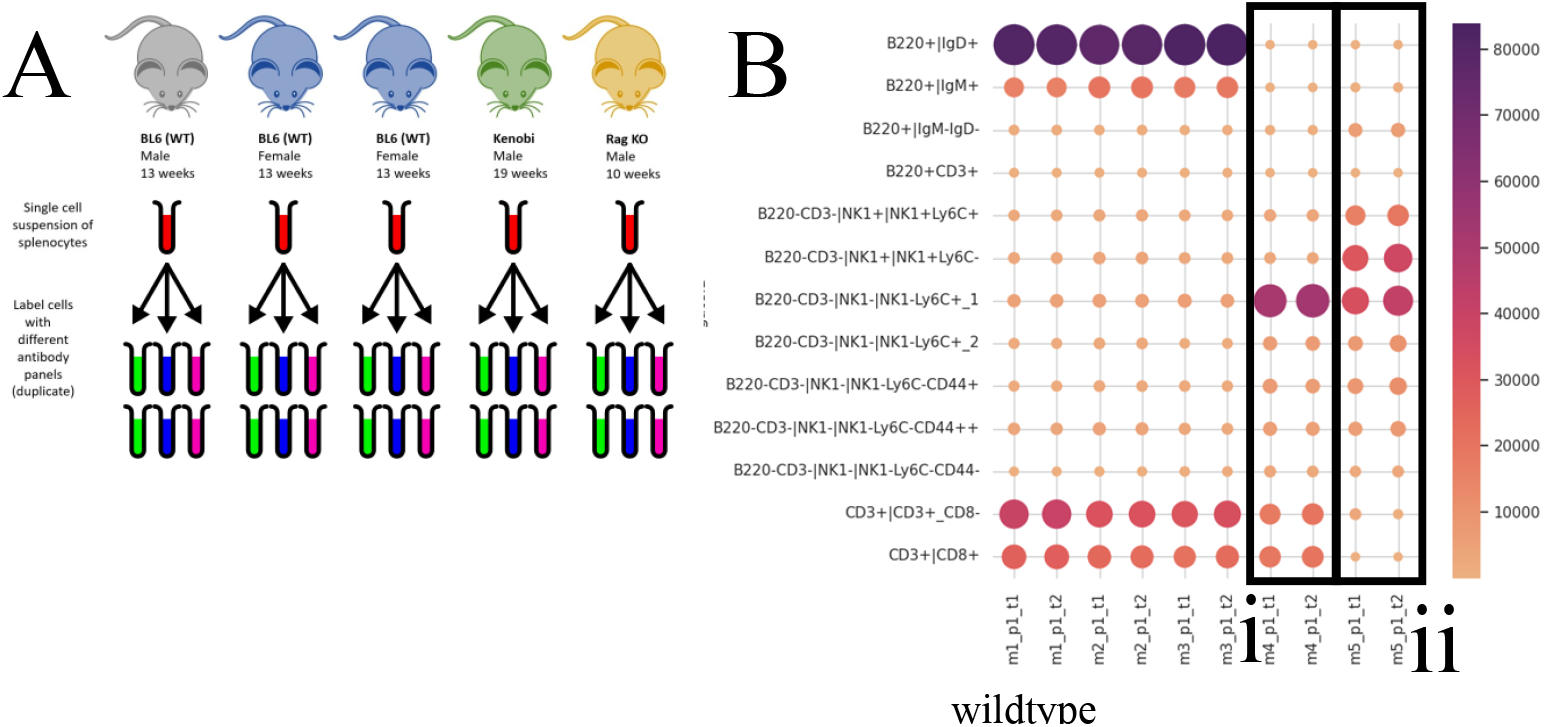
A) Experimental methodology for the synthetic dataset. A single cell suspension of splenocytes was drawn from five mice (three wildtype, one Kenobi, one Rag KO, produced from CRISPR) after 13 weeks. Each suspension was placed on three plates with different antibody panels. Each plate was replicated. B) Heatmap of the cell count for different gates in each sample, with i) Kenobi and ii) Rag KO samples outlined. The area of the circles is proportional to the cell count in each gate.

**Figure 2.**
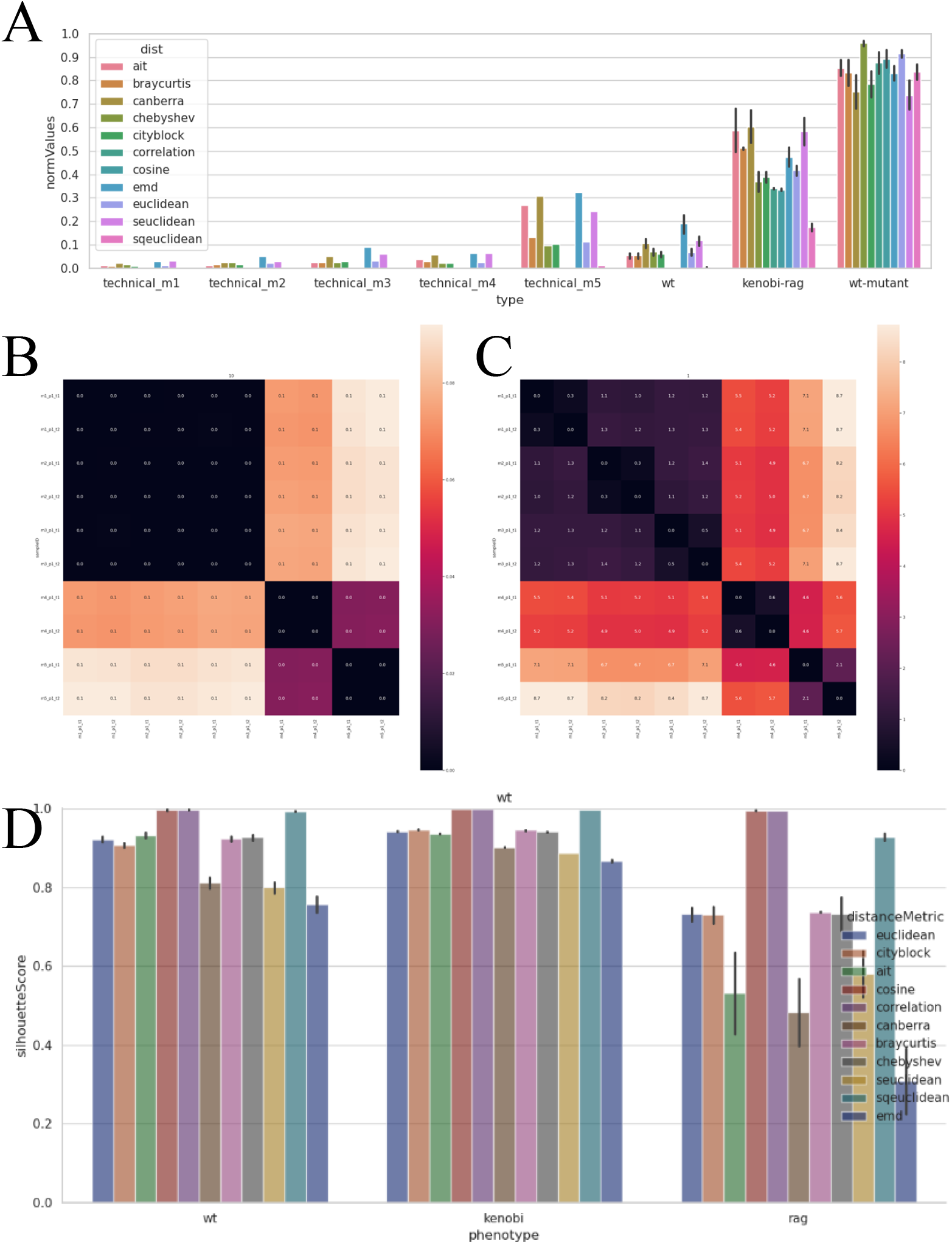
A) Distances of categories grouped by category, normalised to 1. Bars represent mean distance and error bars indicate 95% CI. B, C) Heatmap of the distance matrix between samples in the synthetic dataset. D) Silhouette score of three phenotypes (wildtype, Kenobi, Rag) from the synthetic dataset for each distance metric. Bars represent mean silhouette score, error bars represent standard deviation.

We also observe that Earth Movers Distance (EMD) and standardised Euclidean finds a very high distance between Rag KO’s technical replicates, a high distance between wildtype samples, and a relatively high distance between mutant samples and wildtype-to-mutant samples.

Separation and cohesion were captured with silhouette scores. When observing the technical difference between cluster differentiation, we see that coarse metrics have the highest silhouette scores (Figure 2D), indicating high separation and cohesion.

### Coarse Distance Metrics use Small Cell Populations to Segregate Immunophenotypes

When run on dataset B, cosine similarity clusters appear to be based on gates with relatively small cell counts, as the B220 gates appear not to be of secondary priority (Figure 3; observe sample 18971_Tube_039, which has a very low B220 cell count). Standardised Euclidean appears to largely cluster based on B220 cell count, however not quite capturing sample 18971_Tube_039 in its own cluster as we would expect.

**Figure 3.**
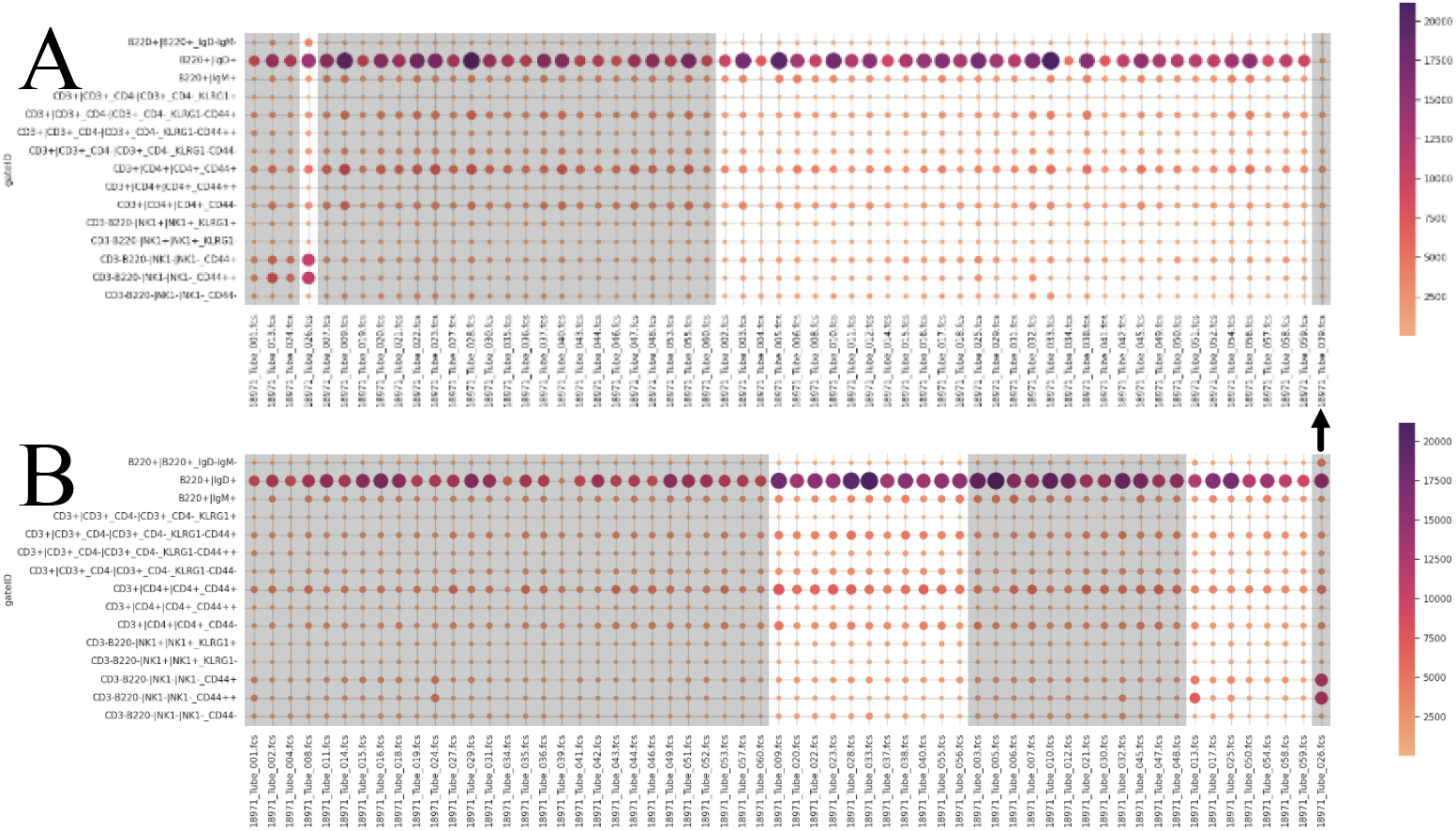
Heatmap of the cell count in each gate for each sample, area size and colour of circles represent cell count of that gate in that sample. Clusters are separated by background colour. A: Cosine similarity, B: standardised Euclidean. The arrow indicates a sample of particular interest.

### Distance Metrics Segregates Mouse Sex and Immunophenotype

Three groups of mutant mice were successfully distinguished from each other and from the large group of wildtype mice by measuring distances between mice with cosine similarity and clustering with hierarchical clustering. The sixty samples in dataset B were split into five clusters. Each cluster aligns with a particular cell phenotype that is observable by Principal Co-ordinates Analysis (PCoA) (Figure 4B). PCoA also demonstrates that the second principal component is mouse sex., demonstrating again that cosine similarity segregates based on biological features. Observing the PCoA plots (Figure 4) and dendrogram (Figure 3) demonstrates that cosine similarity was able to separate mice with clearly similar cell phenotypes.

**Figure 4.**
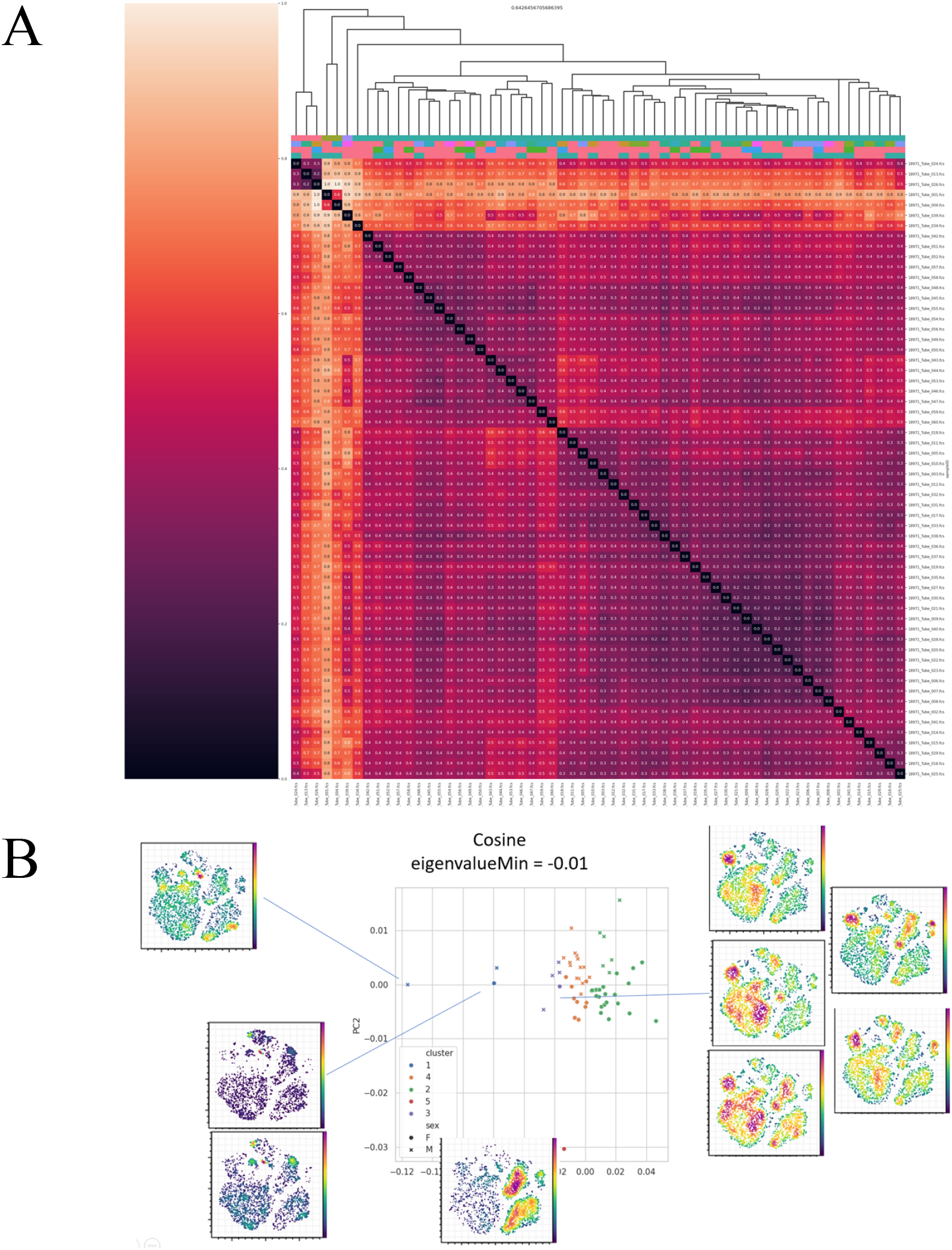
A) Heatmap of the cosine similarity distance matrix for the ENU dataset. The dendrogram was calculated using hierarchical clustering (“maxclust”) and the four colour layers immediately below correspond to: hierarchical clustering clusters, sex, gene of interest, and genotype. B) PCoA plot of the cosine similarity distance matrix for the ENU dataset. Colours refer to hierarchical clustering clusters, marker style refers to sex. tSNE plots are provided for samples of small clusters.

## 3 Discussion

Cosine similarity and standardised Euclidean display qualities that are valuable when classifying flow cytometry data. They are reflective of biological reality, displaying an expected magnitude of distance between wildtype and mutant immunophenotypes. They both produce more separation between different mutant immunophenotypes and cohesion within immunophenotypes than other distance metrics, reflected by their high silhouette scores.

Cosine similarity should be used in non-supervised clustering methods as they are robust to noise, while standardised Euclidean could be used as input for machine-learning models as the noise can be accounted for by the model.

Knowing which distance metric is most suited to the task of analysing flow cytometry data is important. Non-supervised clustering can provide a list of candidate phenotypes to explore, ranked by estimated pathogenicity. However, despite the largely interchangeable nature of distance metrics, the use of the wrong distance metric for the task may produce spurious or inaccurate findings (10). Euclidean distance is most often used in flow cytometry analysis (13, 14), despite theoretical limitations when applied in high-dimensional space (15, 16).

Cosine similarity produces more separated and cohesive clusters than standardised Euclidean. Reflecting biological difference is the most important aspect of determining a distance metric, but consideration must be given to downstream analysis. Distance metrics that produce more separated and cohesive clusters are more useful than those that do not, provided that biological difference is still respected. Clear separation and cohesion of clusters allows for more clear and reproducible downstream analysis.

Observing a heatmap of cosine similarity shows a very clean and clear distinction based on phenotype (Figure 2B,C), whereas a heatmap of standardised Euclidean shows high granularity, identifying differences between technical replicates taken from the same mouse, and identifying that the Rag KO technical replicates are quite different to each other in comparison to other technical replicates. Cosine similarity is useful for the identification of mutant phenotypes while ignoring the noise of variation between individuals that otherwise share the same immunophenotype. Standardised Euclidean is sensitive to this noise, which might be useful as input for sophisticated machine-learning models that can identify the signal in this noise.

Cosine similarity clusters not only by cell populations that are expected to be large, such as cells that express B220 (B cells), but clusters also by cell populations that are, by nature, sparse. Figure 3 shows that cells that were not identified by the eight cell markers used, which typically are quite miniscule in population size, were used to segregate three samples into one cluster, and one sample into its own sample, likely based on the abnormal overexpression of these cells. Similarly, CD4+ cells (helper T cells) are not as plentiful in population size as B cells but were still used to cluster many samples into their own cluster. Standardised Euclidean shares this feature, but notably those clusters with low helper T cells also featured low B cells, indicating that perhaps standardised Euclidean may be clustering simply based on each sample’s total cell population; a useful metric but one that can be calculated without distance metrics.

Mouse sex can be segregated using random forest and flow cytometry immunophenotypes where each marker is split into 400 bins (Dr Benjamin Mashford, personal communication, 2024). We find that cosine similarity and hierarchical clustering alone can cluster mice by sex, further indicating this distance metric is clearly identifying a biological signature, not instrument noise.

Cosine similarity is an informal metric due to violating the triangle inequality property. This is argued to mean that it is unable to be used to rank samples based on their difference from a control (ie wildtype) (11, 17). While a valid formal concern, we have demonstrated that metrics like cosine similarity are capable of distinguishing between mutant samples and wildtype sample, and distinguishing between mutant samples of different phenotypes. Thus, while the difference between mutant samples of different phenotypes may be more difficult to interpret, the high silhouette score and clear cohesion and separation that cosine similarity produces is extremely useful when clustering.

Using flow cytometry data, which is high-dimensional and highly-sensitive, we identified distance metrics that were able to distinguish wildtype from mutant phenotypes without any prior assumptions or labels. These were validated using a synthetic dataset where immunophenotypes were known, and it was shown that coarse distance metrics, cosine similarity, correlation, and squared Euclidean distance, were largely interchangeable for comparing cell populations between samples. These should be used in future experiments as the initial distance metric when comparing FACS immunophenotype data. Standardised Euclidean is most likely to be most useful as input for ML training.

## 4 Methods

All scripts were written and run in Python 3.12.3 and R 4.0 and are hosted on Zenodo (https://doi.org/10.5281/zenodo.14549980). Analysis was conducted on FCS files and associated metadata. Two datasets were used, the first was a synthetic dataset using twelve cell markers (B220, CD3, CD8, CD19, CD25, CD44, CD62L, IgD, IgM, Ly6C, NK1, and LD), the second was an ENU dataset using eight cell markers (B220, CD3, CD4, CD44, IgD, IgM, KLRG1, and NK1).

### ENU Mouse Mutagenesis and Flow Cytometry Preparation

Mouse strains were produced by ENU mutagenesis on a C57BL/6 background at the Australian Phenomics Facility of the Australian National University as described (18). The Kenobi strain was produced as described (19), the Rag1 KO strain was produced as described (20).

Mouse spleen was collected into 3 mL FACS buffer and mashed through a 70 µm cell strainer. Cells were transferred into 15 mL falcon tube, centrifuged at 465 xg, 5 min, 4°C, and supernatant discarded. Red blood cells were lysed by resuspending the pellet in 3 mL 1X lysis buffer (ThermoFisher Scientific, 00-4300-54), incubated for 1 min at room temperature. Post incubation, lysis buffer was diluted by the addition of 7 mL of FACS buffer (2.5% FBS, 0.1% sodium azide, 0.01% EDTA, 10% PBS) and centrifuged at 465 xg, 5 min, 4°C. Supernatant was discarded and the cells were washed by resuspending in 10 mL FACS buffer before being centrifuged at 465 xg for 5 min at 4°C. The cell pellet was resuspended in 0.5 mL FACS buffer and the total cell count was calculated using the Luna-II Automated Cell Counter (ThermoFisher Scientific). Cells were plated into a 96 well round bottom plate.

Cells were blocked in 25 µl of 2X Fc block (BD, Ms CD16/CD32 Pure 2.4G2, 553142) for 5 min before the addition of 25 µl 2X Live Dead stain (ThermoFisher Scientific, Fixable viability dye E780, 65-0865-14). After staining, cells were washed in 200 µl 1xPBS and centrifuged at 465 xg for 5 min at 4°C. Cells were stained with 50 µl of antibody cocktail (in brilliant stain buffer BD Biosciences, 566349) by panels diluted to different antibody concentrations (Table 1) or the respective single colour control at 4°C for 30 min before washing in FACS buffer. Cells were fixed using eBioscience Fix buffer 00-5523-00 according to manufacturer’s instructions. Post fixing, cells were washed twice with FACS buffer before resuspending in 80 µl of FACS buffer for acquisition on the LSRFortessa^™^ X-20 (BD) at the Cytometry, Histology and Advanced Spatial Multiomics (CHASM) Facility, JCSMR.

**Table 1.** Dilution factors for each panel and marker-fluorophore combination. ie 1 in 200. Each antibody is described by its marker (the antigen it binds to on the cell), the fluorophore, the dilution factor of that antibody in that panel, and the catalogue and supplier of the fluorophore. Panel 1 is the optimal antibody concentrations determined by antibody titration.

| Marker | Fluorophore | Panel 1 | Panel 2 | Panel 3 | Catalogue | Supplier |
| --- | --- | --- | --- | --- | --- | --- |
| CD3 | FITC | 200 | 200 | 300 | 100204 | Biolegend |
| CD25 | PE | 400 | 400 | 400 | 102008 | Biolegend |
| CD8 | CD8 | 100 | 400 | 400 | 563786 | BD |
| CD44 | PacBlue | 100 | 400 | 400 | 103020 | Biolegend |
| CD62L | BV605 | 400 | 600 | 600 | 104438 | Biolegend |
| CD19 | BV510 | 600 | 800 | 800 | 115546 | Biolegend |
| IgD | PerCPCy5.5 | 200 | 800 | 800 | 564273 | BD |
| IgM | R718 | 100 | 100 | 600 | 752171 | BD |
| NK1.1 | APC | 200 | 400 | 600 | 550627 | BD |
| Ly6C | PECy7 | 200 | 200 | 400 | 128018 | Biolegend |
| B220 | BUV737 | 300 | 600 | 800 | 612838 | BD |
| CD4 | AF700 | 200 | 300 | 600 | 100430 | Biolegend |
| LiveDead | ef780 | 400 | 600 | 800 | 65-0865-14 | Thermofisher |

### Clean-up and Compensation by Manual Gating

All analysis of FCS files was conducted with CytoExploreR 2.0.0 (21). FCS files underwent logicle transformation to represent the data more appropriately than logarithmic transformation (22). We then removed cellular debris and non-single-cell fluorescence events by gating to isolate events that represented only single cells.

Compensation FCS files were cleaned up according to the above method. They were used to perform create the spillover matrix for the experimental FCS files using the Bagwell-Adams method (23). Experimental FCS files were then cleaned up. The spillover matrices were then used to compensate the experimental FCS files using FlowKit 1.1.1.

### Producing Cell Count Vectors by Manual Gating

After clean-up and compensation, plates were run through dimensionality reduction using FI-tSNE (24) (after down-sampling to accommodate hardware limitations) instead of PCA as non-linear dimensionality reduction provides improved resolution (25). Visual assessment of the hierarchical relationship between cell markers revealed by tSNE plots for one plate in the synthetic dataset and ENU dataset yielded common patterns which were codified in the assignment scheme (Figure 5, Figure 6), segregating each sample into cell populations. Cells were then manually assigned according to this assignment scheme through manual gating.

**Figure 5.**
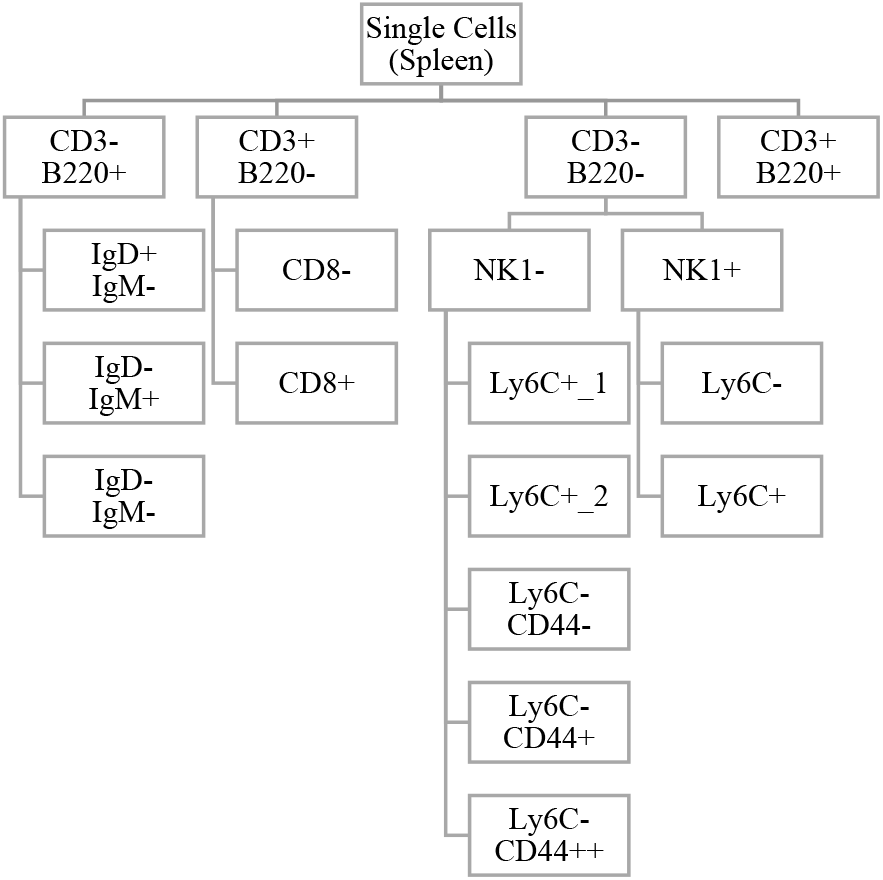
Cell type hierarchy for the synthetic dataset as determined by visual assessment of t-SNE plots. In this assignment scheme, -: cell marker is not/minimally expressed; +: cell marker is moderately expressed; ++: cell marker is highly expressed.

**Figure 6.**
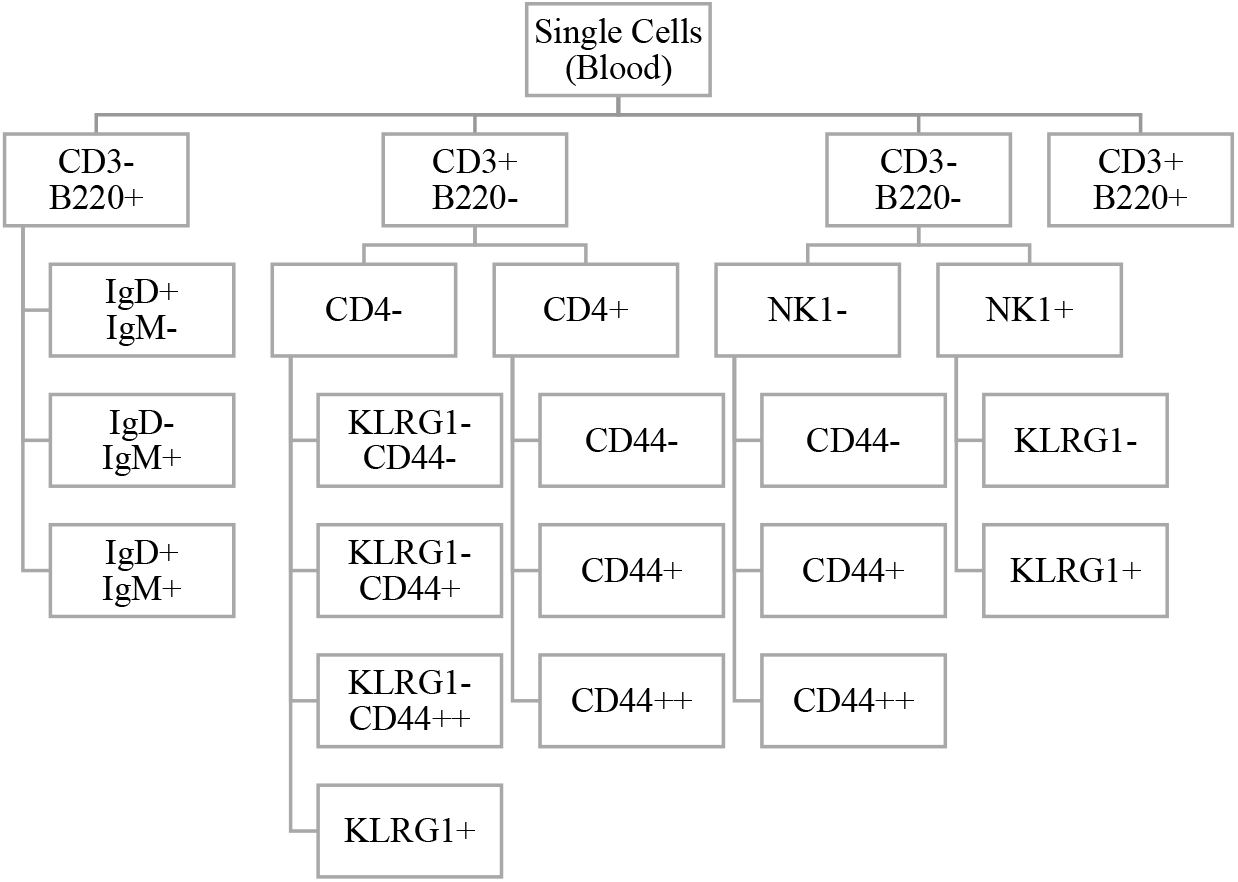
Cell type hierarchy for the ENU dataset as determined by visual assessment of t-SNE plots. In this assignment scheme, -: cell marker is not/minimally expressed; +: cell marker is moderately expressed; ++: cell marker is highly expressed.

### Distance Metrics and Hierarchical Clustering Analysis

Distance matrices were produced using scipy 1.13.0 for the following metrics: Euclidean, Manhattan, Aitchison, cosine similarity, Pearson correlation coefficient, Mahalanobis, Canberra, Braycurtis, Chebyshev, standardised Euclidean, and squared Euclidean. The cell populations in each sample was used as the input, producing a matrix for the distance between each sample. This matrix was then used as input for HCA (Hierarchical Cluster Analysis) using scipy. Linkage was calculated using the “ward” method, and clusters calculated using the “maxclust” method.

When using Aitchison distance, I used multiplicative simple replacement (26) from scikit-bio 0.6.0 to account for zeroes.

### Comparison of Distance Metric Outputs

Silhouette scores were calculated using scikit-learn 1.4.2.

## Notes

### Competing Interest Statement

The authors have declared no competing interest.

